# Fluid-structure Interaction Simulation of the Cerebrovascular Circulation: Immersed Boundary versus Arbitrary Lagrangian Eulerian Mesh Formulations

**DOI:** 10.1101/2025.09.28.679107

**Authors:** Maximos P McCune, Daniel E Thomeer, Mark A Davison, Majid Rashidi, Nina Z Moore

**Affiliations:** Cerebrovascular Department, Neurologic Institute, Cleveland Clinic Foundation, Cleveland, OH, USA; Department of Neurosurgery, Neurologic Institute, Cleveland Clinic Foundation, Cleveland, OH, USA; Department of Biomedical Engineering, Lerner Research Institute, Cleveland Clinic Foundation, Cleveland, OH, USA; Cleveland Clinic Lerner College of Medicine, Cleveland Clinic Foundation, Cleveland, OH, USA

**Keywords:** Fluid-structure interaction, Immersed boundary simulation, Arbitrary Lagrangian Eulerian simulation, Cerebrovascular modeling, Internal carotid artery

## Abstract

**Purpose:** Patient-specific fluid-structure interaction (FSI) simulations allow for the *in silico* modeling of vascular pathology. Though existing attempts to model the cerebrovasculature confirm the potential of FSI as a future diagnostic tool, standard simulation methods and modeling parameters remain undefined. The purpose of this investigation was to compare immersed boundary (IB) and arbitrary Lagrangian Eulerian (ALE) formulations to discern whether the increased modeling complexity offered by ALE is necessary for the modeling of smaller-caliber vessels, given increased computational load.

**Methods:** Direct comparisons of Fluent and Mechanical behavior were conducted between IB and ALE methods of FSI simulation. Simulations utilized an internal carotid artery geometry conduit with optimized mesh. Boundary constraints were derived from previous investigation of vascular tissue and fit to a Prony series. Both qualitative profile comparisons and quantitative parametric analyses of variance were conducted to assess differences in simulation output.

**Results:** In this study, we report deviations in Fluent and Mechanical output between IB and ALE cases of FSI simulation. More specifically, ALE-method simulations boast higher stress, lower wall shear stress, and lower strain. These differences persist across the vessel geometry and increase with high strain. Additionally, inconsistencies between solving methods are exacerbated in areas of more complex mesh geometry (i.e. vessel bifurcation).

**Conclusion:** Substantial alterations in intraluminal stress, shear stress, and strain suggest that ALE formulation is necessary for modeling blood vessels of the cerebrovasculature. Our findings highlight the importance of accurately modeling the dynamic interactions that occur between the fluid and material domains of simulation.

## INTRODUCTION

Fluid-structure interaction (FSI) is a computational method that simulates the interplay between a fluid and solid at the phase interface. FSI is useful for providing accurate modeling of systems that require meticulous consideration of both the mechanical and fluid domains; such is the case in the computational modeling of blood flow through angioarchitecture. Establishing image-based hemodynamic models offers new ways to study and predict the outcomes of vascular disease. Vascular FSI research aims to accurately simulate hemodynamics and associated mechanical deformations in the vessel walls to facilitate the design of targeted endovascular therapies and further our understanding of vascular pathology [1,2]. As such, ensuring modeling accuracy of FSI simulations is paramount if future simulations are to guide clinical decisions and act as a qualifying metric for surgical intervention.

*In silico* trials conducted by Moerman et al. serve to demonstrate the feasibility of patient-specific FSI modeling in cerebrovascular study, succeeding in the generation of an arbitrary Lagrangian Eulerian (ALE)-like [3], finite elements simulation of a carotid bifurcation using FEBio software [4]. Similarly robust models have been constructed for the study of cerebral aneurysms to assess rupture risk and the application of endovascular flow diverter placement and coiling [5 and 6]. Valencia et al. reported marked differences in wall shear stress (WSS) across cerebral aneurysm morphologies with a fully coupled fluid-structure model of patient vessels, and Torii et al. found similar success with a model of aneurysm hyperelasticity [7 and 8]. Though FSI modeling shows clear promise as a guide for vascular intervention, precise simulations have yet to be devised for the study of more intricate vascular pathologies, such as arteriovenous malformation [2]. Such computationally demanding simulations, however, present an issue of solving efficiency—requiring further optimization.

To ease computational load, many initial approximations of blood flow involved fixed, rigid vessels and assumed zero wall motion [9]; although, such simplifications likely mar the efficacy of a model intended for translational use with clinical relevance. For this reason, selecting a set of assumptions that adequately balances processing time and modeling accuracy is crucial. The primary concern of this study was to contrast specificity and efficiency between monolithic and partitioned solving methods for the simulation of blood flow through cerebrovascular tissue. While the non-conforming mesh of a monolithic approach boasts faster solving speeds, it lacks the rigorousness gleaned from the two-way convergence of partitioned FSI [10]. An adequate simulation of mechanical behavior is necessary for accurate simulation—as wall rigidity has been noted to impact simulated realism of stress on vascular tissue [11]—but the degree to which a more complex formulation relevantly enhances a model of small-caliber vessels is unknown. As such, we considered whether the immersed boundary (IB) formulation or ALE finite element method is better suited to discretize the fluid and structure boundary domains for simulations of flow through a vessel.

With the IB method, structural domains are modeled with a Lagrangian mesh and allowed to move freely against a Eulerian fluid. Since no re-interfacing occurs between the solid and fluid elements, this process is well-suited for modeling systems with predictable variance in the structural domain; however, the IB method lacks a rigorous description of fluid behavior. The ALE method, while more computationally intensive and prone to technical restraints, involves simultaneous reformulation of both the structural and fluid domains to offer a higher degree of modeling accuracy [12]. The fluid domain is modeled as a negative solid and responds accordingly to changes in the surrounding structure at the material interface. At each step, the velocity at the structural grid is solved using pseudo-elasticity equations and translated to the fluid— providing a more detailed representation of the fluid-structure interactions with constant, reiterated solving of said fluid. As such, the ALE method is better equipped to proxy systems with irregular, moving boundaries. To assess whether it is necessary to incur the additional computational time required of the ALE method for the modeling cerebral blood vessels, we compared flow parameters and stress-strain interactions in multipulse FSI simulations of blood flow between the IB and ALE methods, utilizing a human internal carotid artery model as the conduit.

## MATERIALS AND METHODS

### I. Hardware Specifications

The implementations of FSI simulation in this study were carried out on a single machine utilizing 16 cores for ANSYS solvers. Hardware included 2 Intel Xeon Silver 4210R 2.4GHz, (3.2 GHz Turbo, 10C, 9.6 GT/s 2 UPI, 13.75 MB Cache) processors and a Nvidia RTX A4000 (16 GB, 4 DP) GPU. All simulations were run locally, without the use of a cluster.

### II. Geometric Modeling and Mesh Construction

FSI simulations were performed using ANSYS Fluent and Mechanical software version 2023 R2 (ANSYS Inc., Lebanon, NH). The left internal carotid artery (ICA) bifurcation geometry was modeled from patient angiography using Mimics Medical version 23.0 and 3-matic version 15.0 software (Materialise NV, Leuven, Belgium) and transferred as a STL file into ANSYS workbench. The vessel wall was constructed to a thickness of 0.6 mm with ANSYS Designer software version 2023 R2 (ANSYS Inc., Lebanon, NH), and a 25 mm entrance length was added onto the vessel inlet to establish a fully developed flow profile within the geometry of interest (**Fig. 1**). For Fluent simulations, a tetrahedron mesh was created with an element size of 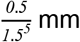, while Mechanical simulations, used a mesh element size of 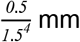 . In both cases, the mesh size of the entrance length was set to 1 mm to decrease simulation time, as no data was recorded in this region. Mesh sizes of relevant geometries were determined through a mesh convergence study starting at 0.5 mm and decreasing sequentially by an element size factor of 1.5^*n*^ until the solution converged (**Table 1**). Over the course of both IB and ALE simulations, no remeshing occurred in either the Mechanical or Fluent domain. Wall motion was minimal, such that no additional nodes were added during modeled flow.

**Table 1:**
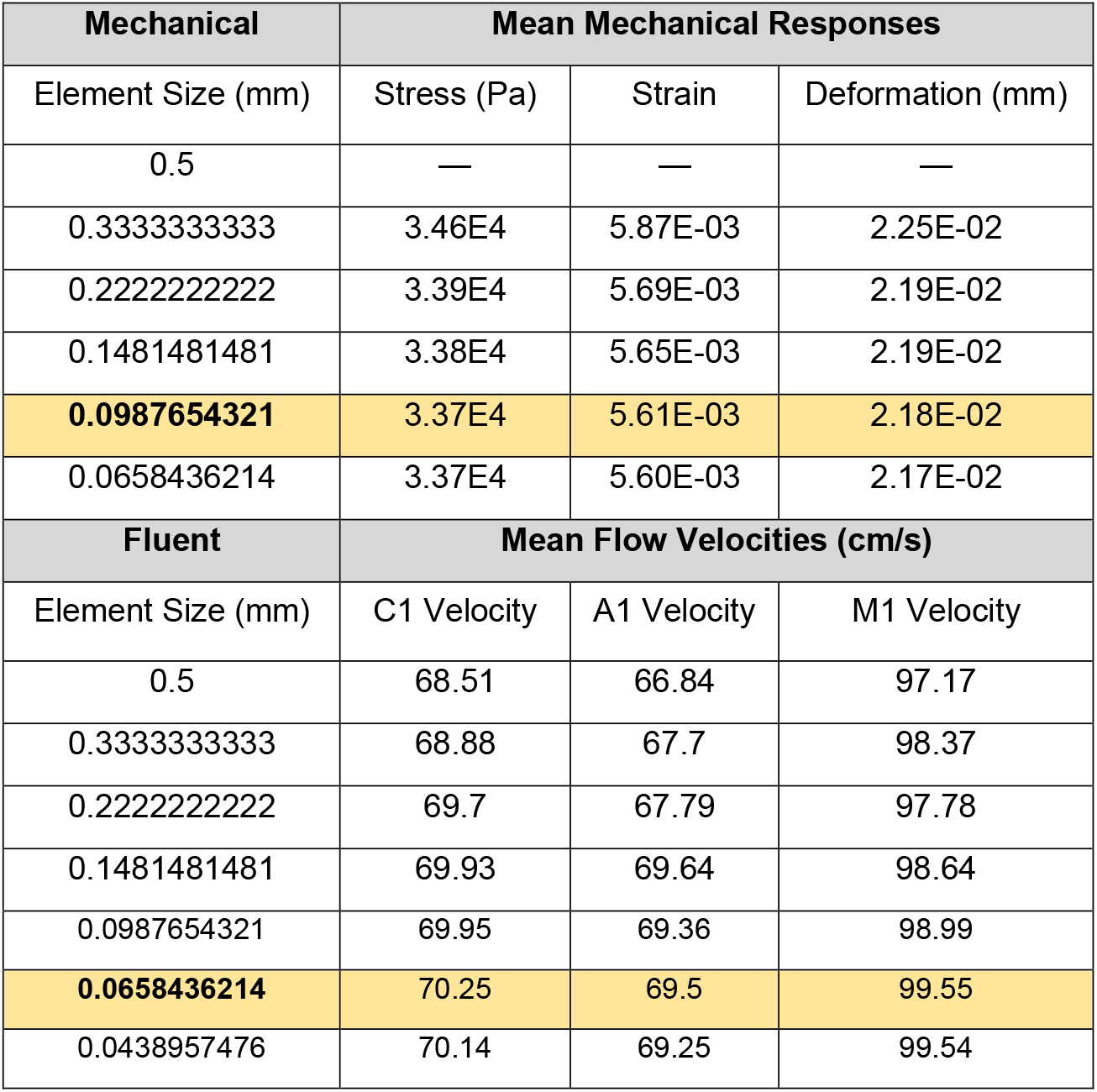
Mesh convergence study results for both Mechanical and Fluent elements. Bolded values signify mesh sizes used in this study. All meshes were constructed in sequence with the formula: 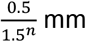, where *n* = study iteration.

| Mechanical | Mean Mechanical Responses |  |  |
| --- | --- | --- | --- |
| Element Size (mm) | Stress (Pa) | Strain | Deformation (mm) |
| 0.5 | — | — | — |
| 0.3333333333 | 3.46E4 | 5.87E-03 | 2.25E-02 |
| 0.2222222222 | 3.39E4 | 5.69E-03 | 2.19E-02 |
| 0.1481481481 | 3.38E4 | 5.65E-03 | 2.19E-02 |
| **0.0987654321** | 3.37E4 | 5.61E-03 | 2.18E-02 |
| 0.0658436214 | 3.37E4 | 5.60E-03 | 2.17E-02 |
| Fluent | Mean Flow Velocities (cm/s) |  |  |
| Element Size (mm) | C1 Velocity | A1 Velocity | M1 Velocity |
| 0.5 | 68.51 | 66.84 | 97.17 |
| 0.3333333333 | 68.88 | 67.7 | 98.37 |
| 0.2222222222 | 69.7 | 67.79 | 97.78 |
| 0.1481481481 | 69.93 | 69.64 | 98.64 |
| 0.0987654321 | 69.95 | 69.36 | 98.99 |
| **0.0658436214** | 70.25 | 69.5 | 99.55 |
| 0.0438957476 | 70.14 | 69.25 | 99.54 |

**Fig. 1:**
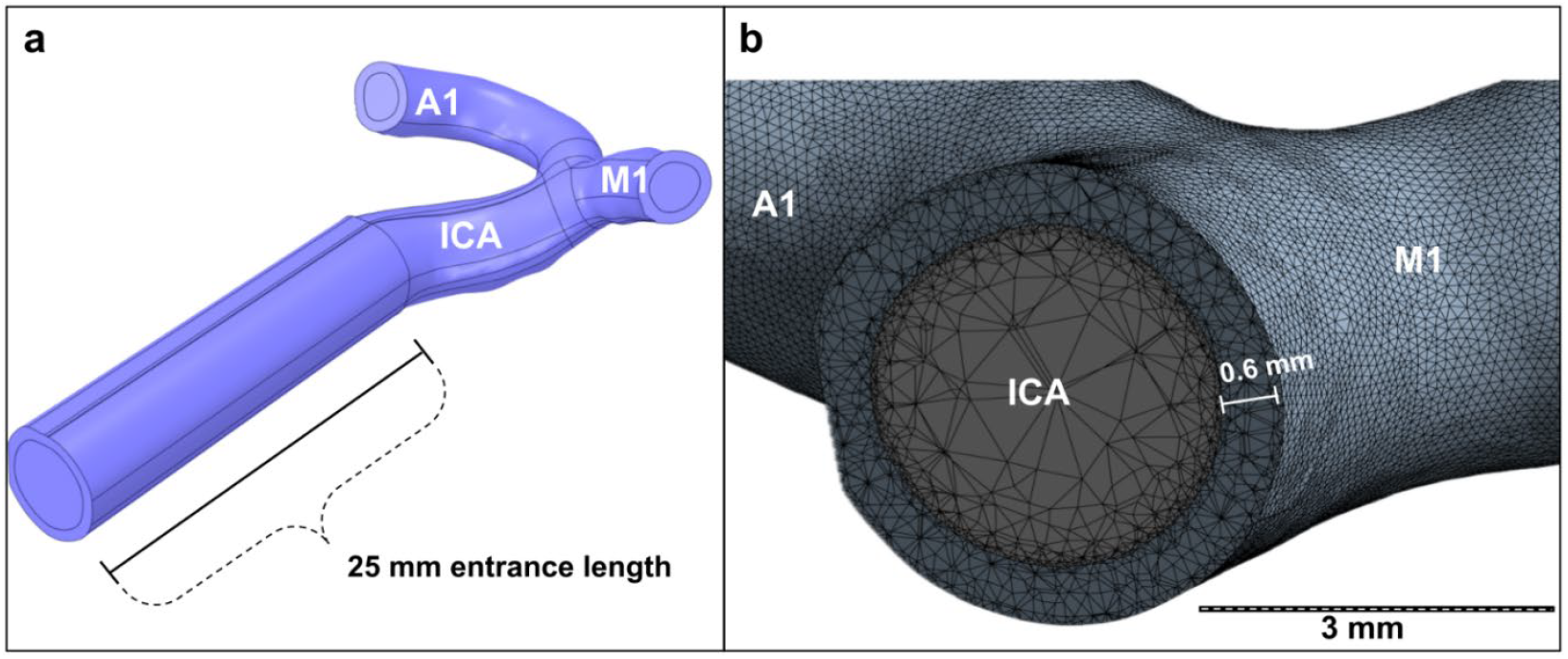
Simulation geometry used in this study (a). Mesh of Fluent (gray) and Mechanical (blue) elements used for the simulation comparison of IB and ALE methods (b). The longitudinal slice depicts the Fluent-Mechanical mesh interface distal to a 25 mm entrance length.

### III. Fluent Simulation

Fluent simulations were carried out using the Semi-Implicit Method for Pressure-Linked Equations (SIMPLE) solver and second order spatial discretization methods. Blood was modeled as a non-Newtonian fluid with the Carreau equation coefficients, time-constant of *2*.*517 s*, power-law index of *0*.*5736*, zero shear viscosity of *0*.*0314* and infinite shear viscosity of *0*.*0037* as seen in prior studies [13]. Blood density was set as *1056* [13]. The blood flow was laminar with a sinusoidal input of the equation 1

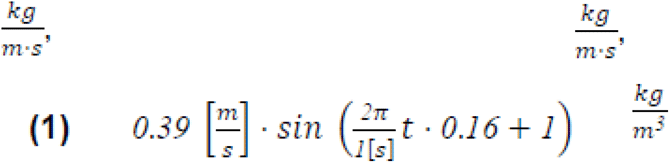

Blood pressure throughout the bifurcation was set to 90 mmHg at the conduit inlet and outlets to match prior *in vivo* measurements of the ICA bifurcation [14].

### IV. Mechanical Simulation

Mechanical properties of the vessel were taken from quantifying Mechanical inflation testing as a Prony series with two terms which has been shown to sufficiently fit viscoelastic data [15]. The Prony series models a linear viscoelastic material by fitting experimental data using the instantaneous modulus (1.87E+7 Pa), Prony constants (0.70827, 0.036246), and Prony retardation time constants (0.58307 s, 5.29E+16 s). The ends of the vessel conduit were fixed in space, and the interior of the vessel was coupled to the Fluent simulation to mimic reactionary forces that would be applied from beyond the segment *in vivo*.

### V. Modeling Parameters

IB method simulations were generated in a stepwise manner, by first solving the simulation in Fluent and then transferring the vessel wall pressure to Mechanical. ALE simulations were solved using system coupling in Workbench to facilitate force transfer from Fluent to Mechanical, communicating wall displacement between Mechanical and Fluent solvers in a simultaneous manner. Both methods were run for a period of four seconds across 400 time steps. Data was recorded from the final two seconds of the simulation to ensure a fully developed flow profile.

### VI. Probe Placement

Measurement probes were placed at specific locations throughout each segment of the vessel to collect velocity and wall shear stress from the Fluent simulations and stress, and strain data from the Mechanical simulation. Velocity probes were positioned in the middle of the ICA, A1 (the proximal-most segment of the anterior cerebral artery), and M1 (the proximal-most segment of the middle cerebral artery) to capture measurements at each branch. Strain, stress and WSS measurements were captured by probes placed circumferentially about the conduit inlet and each outlet. For a single time-step study, probes were placed at the vessel entrance (just distal to the entrance length segment) and around the ICA bifurcation, allowing for direct comparison between mesh structures.

### VII. Simulation Contour Mapping & Statistical Modeling

Simulation output profiles of velocity, strain, stress, and WSS were prepared from data collected at the final time step, after 2 seconds of flow development and 2 seconds of measured flow. Comparisons were then drawn qualitatively by comparing velocity streamlines and contour maps between IB and ALE solutions. Parametric cohort comparisons between IB and ALE solver methods were conducted using R version 4.3.1 (R Foundation, Vienna, Austria). One-way analysis of variance (ANOVA) and Welch’s ANOVA tests were performed where appropriate to account for data of unequal variance. To assess how variations between methods may differ across geometries, T-tests were similarly conducted in R from data generated in ANSYS. All null-hypothesis testing reported in this study considered *p* < 0.05 as significant for statistical comparisons. Data from every probe was considered in our analysis, with no removal of outliers.

## RESULTS

### I. Flow Velocity Simulation Results

We demonstrate that both IB and ALE simulation methods reliably proxy human blood flow conditions. Though mean flow velocities probed within ICA inlet and outlet geometries were higher than prior *in vivo* measurements collected via transcranial Doppler ultrasonography, simulation values still resemble physiologic data (**Table 2**). ALE-case flow velocity values (ICA: 63 cm/s, A1: 66.3 cm/s, and M1: 89.8 cm/s) were lower than IB simulation outputs (ICA: 70.3 cm/s, A1: 69.8 cm/s, and M1: 98.5 cm/s) and closer to *in vivo* averages, and thus boasted a higher degree of modeling accuracy.

**Table 2:**
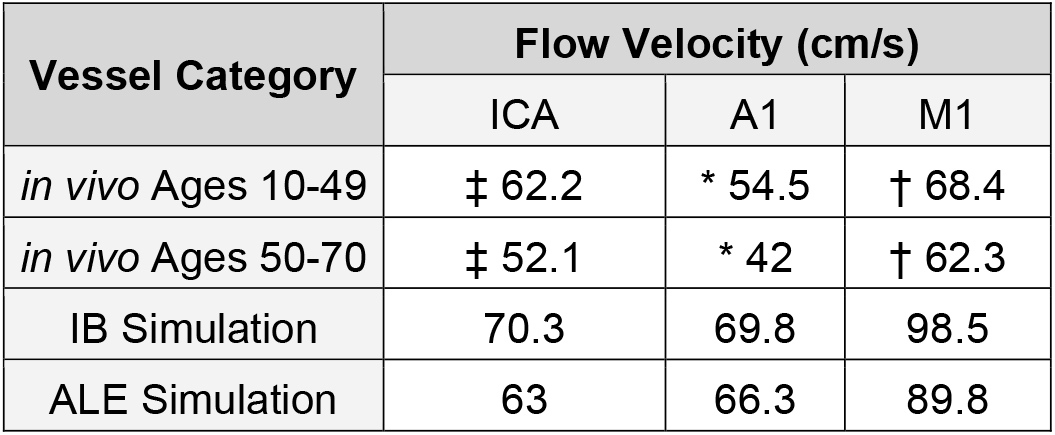
Flow velocities recorded within ICA inlet and outlet geometries, averaged across 2 seconds of simulated flow, compared to in vivo measurements recorded with transcranial Doppler ultrasonography from * Ringelstein et al. 1990, † Nagai et al. 1998, or a combination of ‡ both Ringelstein and Nagai data [26,27].

Visualization of pulsatile flow confirms this reduction in flow velocity in vessel segments solved with ALE formulation when compared to IB solutions under equivalent mesh conditions (**Fig. 2**). These differences in observed flow velocities were consistent across all probed locations, inclusive of ICA, A1 and M1 segments. Velocity streamlines depict the flow discrepancies between ALE and IB methods, with the largest differences measured in areas of highest, most disparate strain (**Fig. 3**). This result is most evident just distal to the bifurcation, where the areas of highest flow are identified to show the greatest strain differences between solver cases. We also note that areas of more convoluted geometry, as seen in the ICA terminus and structures immediately distal, display higher strain and markedly different velocities between methods (**Fig. 3**). These data suggest that ALE iterative solving is most necessary in areas of high tortuosity and elevated vessel wall strain.

**Fig. 2:**
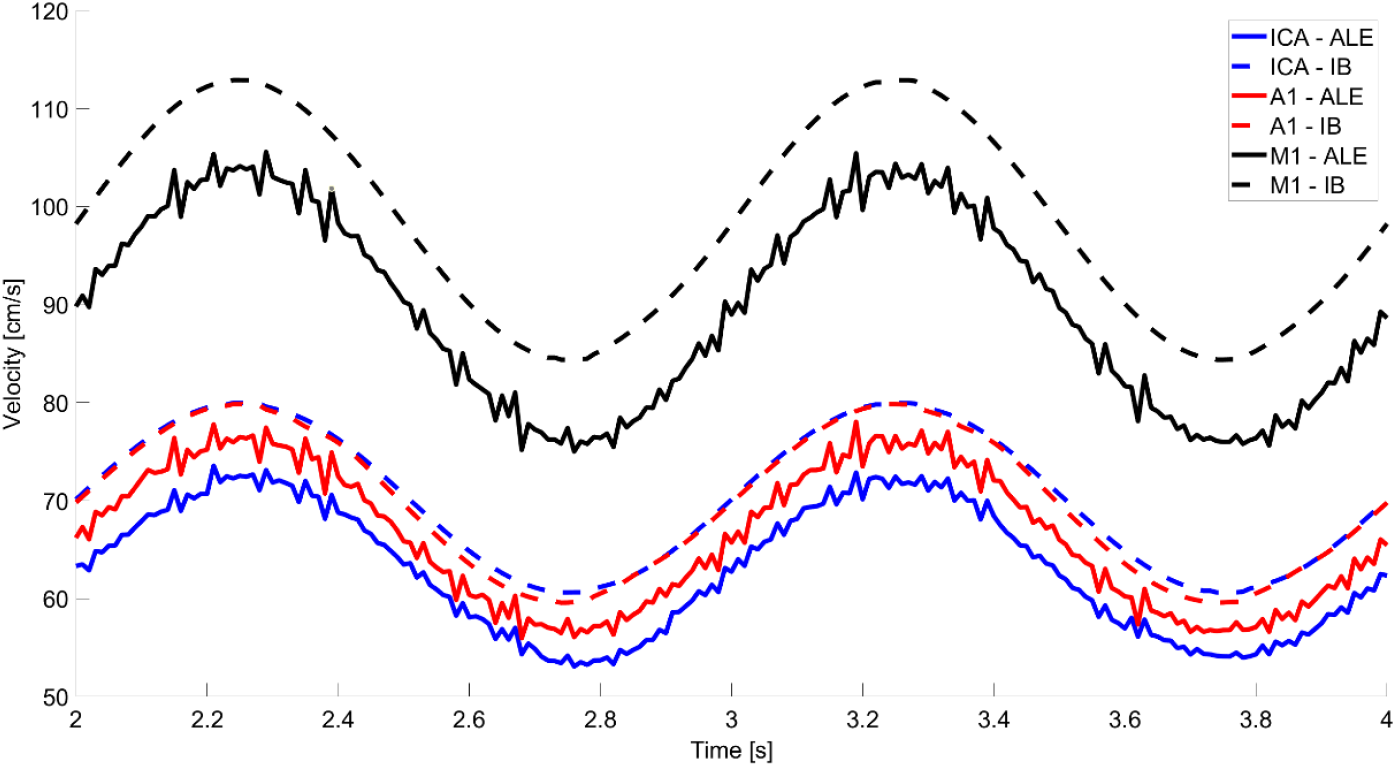
Flow velocity comparison between IB and ALE methods at ICA bifurcation inlet and outlet geometries (A1 and M1).

**Fig. 3:**
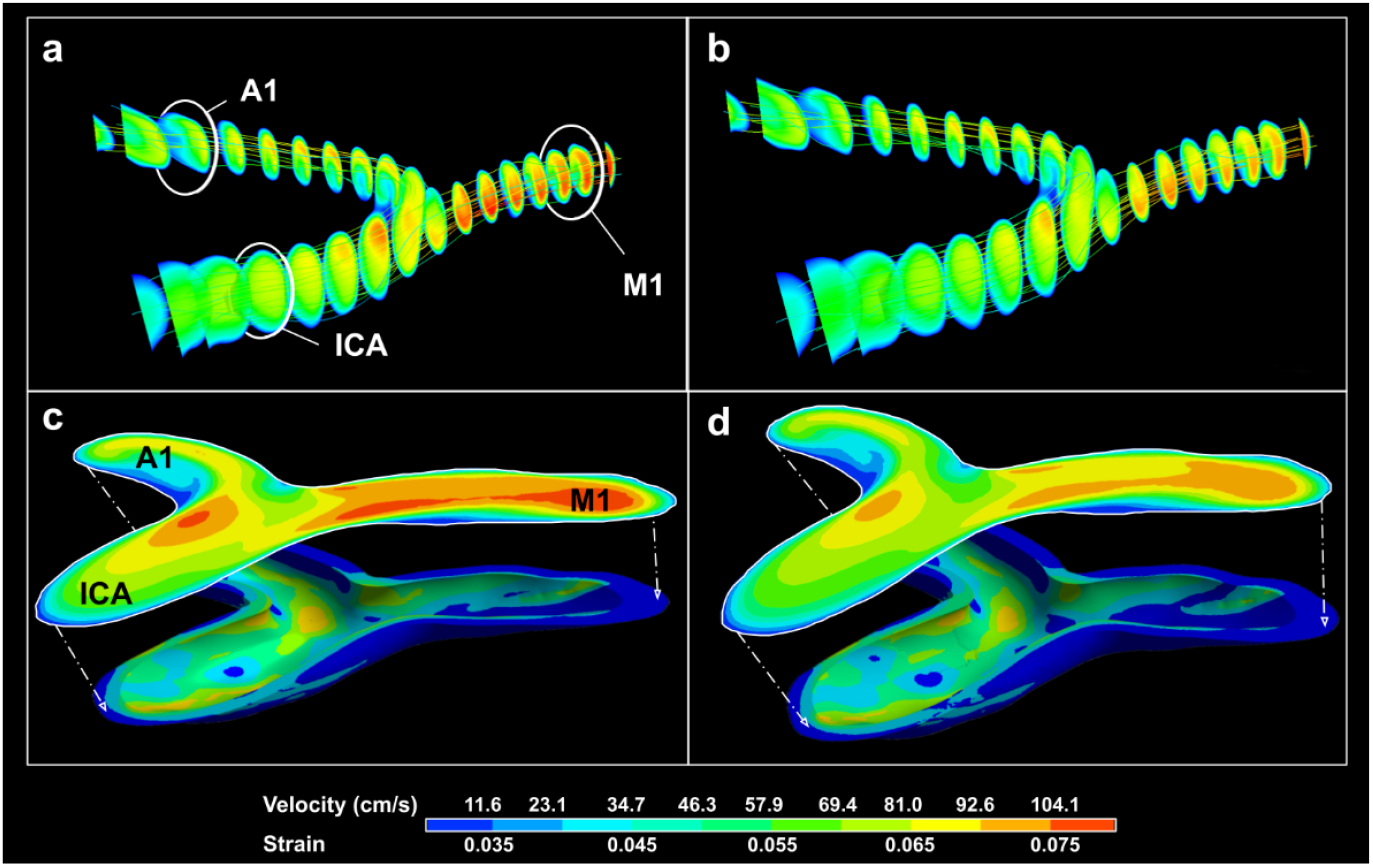
Flow velocity streamlines comparison between IB (a) and ALE (b) methods. Velocities were then displayed against strain in the same regions, with 2-dimensional velocity profiles (upper) overlaid atop strain renderings (lower) between IB (c) and ALE (d) simulations. Streamlines and contour maps were generated from data collected after 2 seconds of simulated flow.

### II. Strain Simulation Results

In an investigation of vessel wall strain, we observed increased measures at all probed locations when using an IB solver (**Fig. 4**), with highest percent strains noted in IB simulations of the M1 inlet territory at 14.18%. IB simulation of the A1 inlet and ICA terminus displayed appreciably lower peak strains at 13.07% and 11.83% respectively, though still higher than all ALE counterparts.

**Fig. 4:**
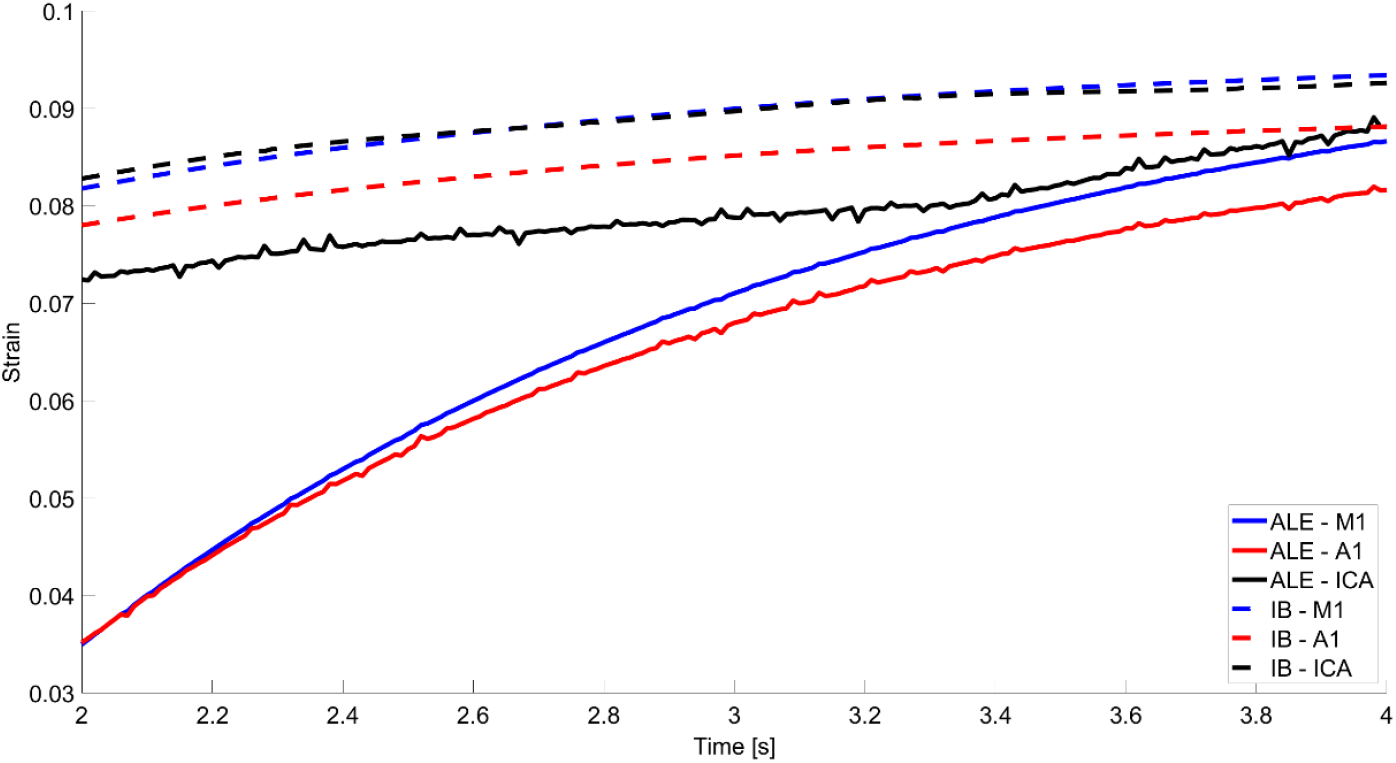
Strain comparison between IB and ALE methods at ICA bifurcation inlet and outlet geometries (A1 and M1).

In addition, we compared strain in a single time-step simulation of flow with a second set of probes confined to the ICA inlet and bifurcation—providing a more robust comparison with consideration for areas of heightened geometric complexity. This second analysis echoes the aforementioned differences between IB and ALE cases (**Table 3**). We recorded significantly lower strains in the ALE case than with a monolithic solver at both the ICA inlet and bifurcation (*p* = 0.03 in both comparisons, **Table 3**), with an average difference between methods of 0.0025 and 0.0027 respectively, and found that strain measurements in IB simulations were more variable (**Table 4**). We report an IB-case mean inlet strain of 0.0316 and mean bifurcation strain of 0.0362, with decreased ALE measurements of 0.0291 mean inlet strain and mean bifurcation strain of 0.0335.

**Table 3:**
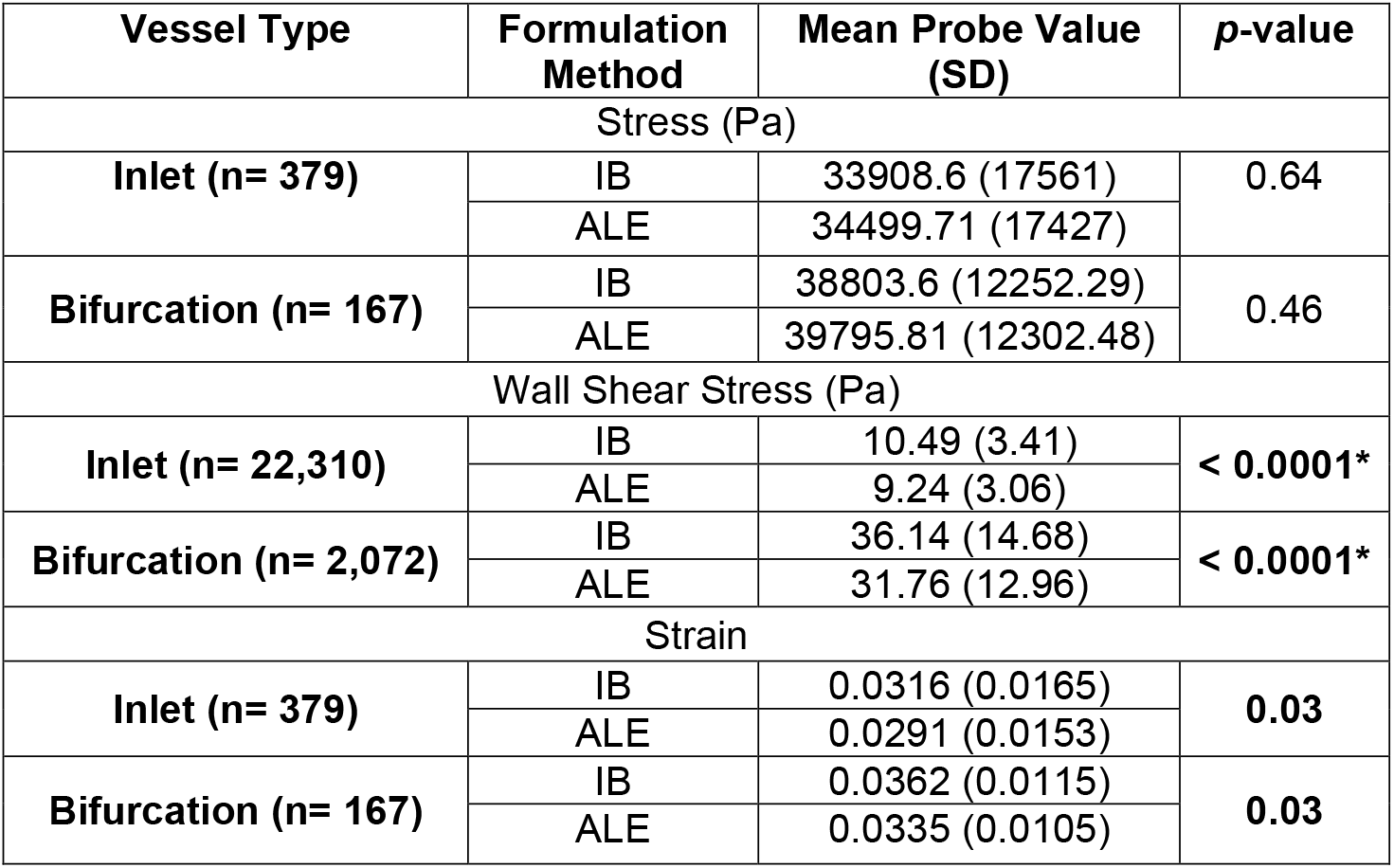
Single time-point comparison of ICA inlet and bifurcation probe values between IB and ALE formulation methods. Values reported are the average across probes after 2 seconds of simulated flow, utilizing one-way ANOVA and Welch’s ANOVA (*) where appropriate.

**Table 4:** Comparison of reported IB-ALE mean difference outcomes between inlet and bifurcation ICA geometries recorded at each probe locus. Student’s and Welch’s t-test (*) were conducted on values after 2 seconds of simulated flow.

| Vessel Type | Mean Difference Between Methods | *p*-value |
| --- | --- | --- |
| Stress (Pa, n= 546) |  |  |
| Inlet | 591.11 | < 0.001* |
| Bifurcation | 992.21 |  |
| Wall Shear Stress (Pa, n= 24,392) |  |  |
| Inlet | 1.25 | < 0.0001* |
| Bifurcation | 4.38 |  |
| Strain (n= 546) |  |  |
| Inlet | 0.0025 | < 0.001 |
| Bifurcation | 0.0027 |  |

Further, this average difference in strain recorded between IB and ALE cases was lower across inlet probes than at the bifurcation (*p* < 0.001), again suggesting that iterative formulation is better suited to model less uniform angioarchitecture (**Table 4**).

Reinforcing this notion with strain profile comparisons (**Fig. 5**), we continued to find that strain is markedly disparate between IB and ALE cases. Our data shows more diffuse regions of high strain despite lower overall strain, which together establishes ALE as the more robust solver for the simulation of the cerebrovasculature and should be prioritized in simulations of a complex mesh with clinical relevance.

**Fig. 5:**
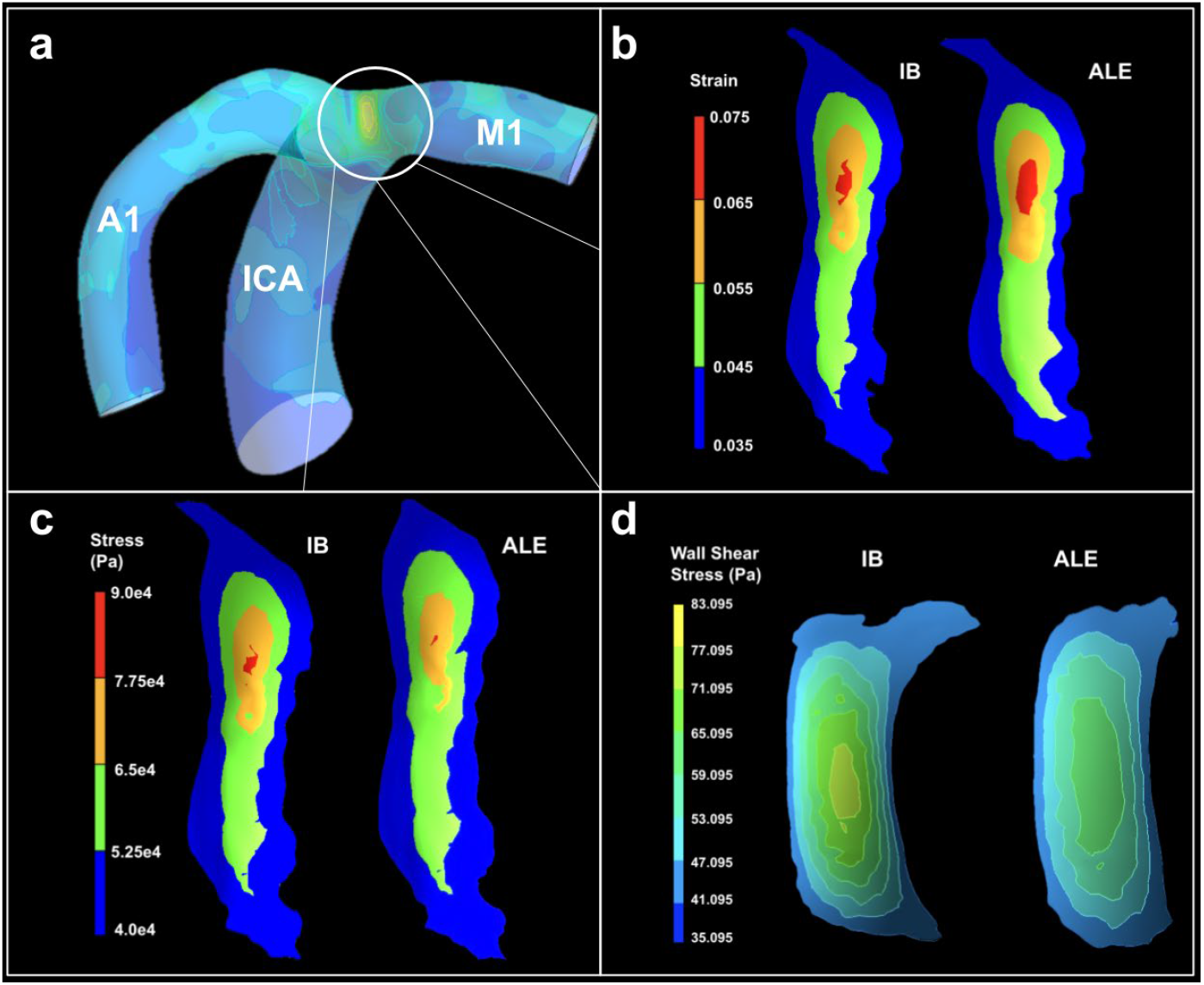
FSI contour mapping comparisons between IB and ALE formulation methods of strain, stress, and WSS after 2 seconds of simulated flow. Full-scale ICA geometry is shown in panel a. Panels b-d depict mechanical profiles at the ICA bifurcation.

### III. Stress and Wall Shear Stress Simulation Results

In an analysis of wall stress, we found that ALE cases displayed higher stresses than IB analogs at all probed inlet and outlet locations (**Fig. 6**). ALE results also showed higher fluctuations in vessel stress, due to the two-way transfer of information between Fluent and Mechanical modules, as seen in the noise captured in Figure 6 with ALE cases. A1 inlet probes recorded the lowest general stress, but IB and ALE differences varied similarly to ICA and M1 probes (**Fig. 6**). We also observed a more pronounced dampening of stress in ALE cases. This reduction in IB stress periodicity is most prominent in ICA measurements, as these probes precede the exacerbated dampening caused by splitting at the bifurcation. Though still perceptible, this effect is diminished in A1 and M1 geometries (**Fig. 6**).

**Fig. 6:**
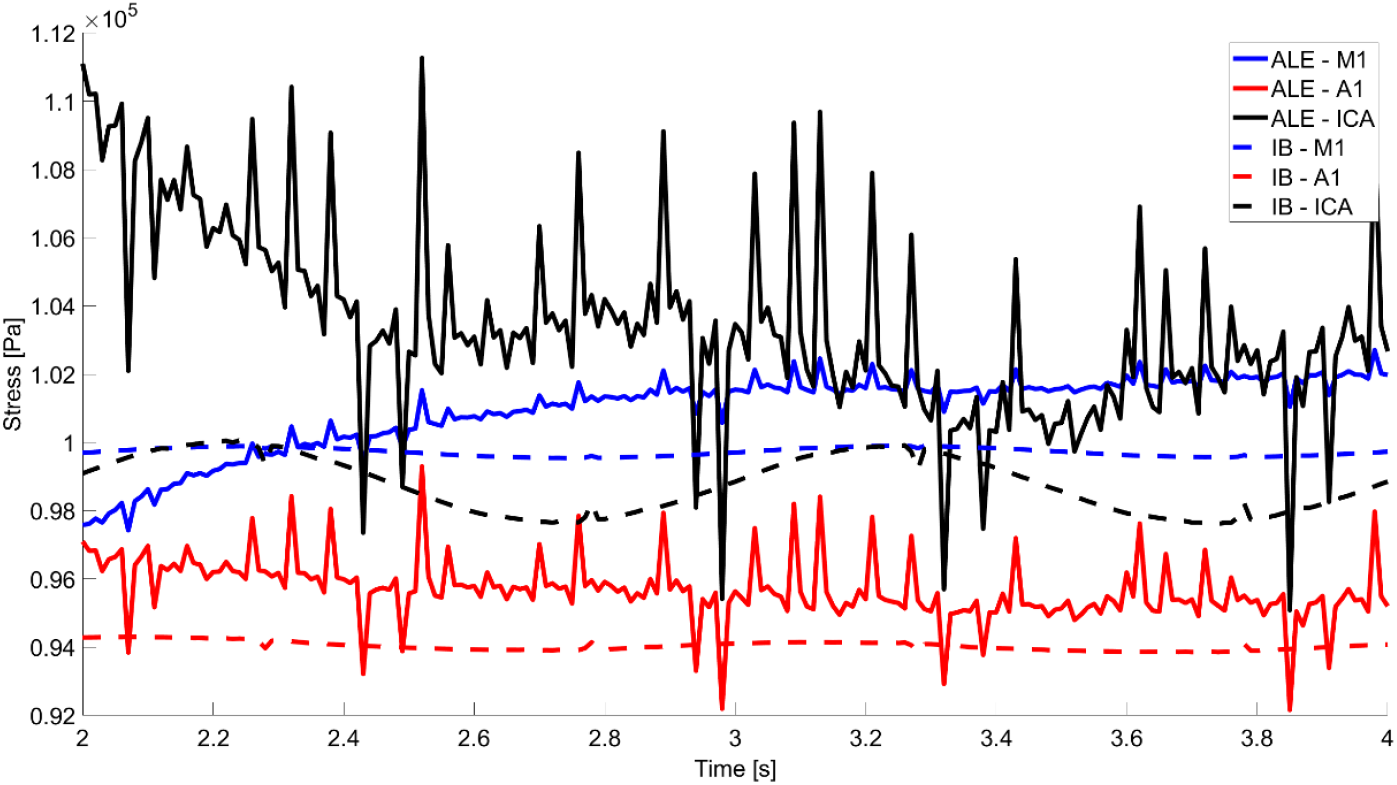
Stress comparison between IB and ALE methods at ICA bifurcation inlet and outlet geometries (A1 and M1).

While not to the degree of statistical significance (*p* = 0.64 at the inlet and *p* = 0.46 at the bifurcation), single time-point comparisons between IB and ALE methods confirm higher stresses with ALE formulation across both probed locations within the simulation structure. Specifically, we report a mean ALE stress of 34499.71 (SD ± 17427) Pa at the ICA inlet with a mean IB stress of only 33908.6 (SD ± 17561) Pa. At the bifurcation, we again found higher mean stress in the ALE case at 39795.81 (SD ± 12302.48) Pa, compared to 38803.6 (SD ± 12252.29) Pa with IB formulation (**Table 3)**.

On further analysis, mean difference comparisons of solver method between inlet and bifurcation probe sets revealed that the magnitude of variance between IB and ALE stress measures significantly differs between locations about the vessel interior, suggesting a rationale for using ALE formulation to simulate increasingly convoluted geometries. Our study revealed an average difference between methods at the inlet of 591.11 Pa and an average difference between methods at the bifurcation of 992.21 Pa (*p* < 0.001, **Table 4**). Stress profiles at the ICA bifurcation offer additional support for this nuanced difference, as stress is generally higher but more diffuse in ALE simulations (**Figs. 5 and 6**).

Regarding WSS, we again demonstrate differences between IB and ALE methods (**Fig. 7**). WSS measures, consistent across ICA, A1, and M1 inlet locations, were lower in ALE cases. As expected, these findings mirror the decreases in flow velocity found with ALE formulation. Diminished flow velocity reduces shearing across the vessel wall interface, lending to the decrease in WSS. Bifurcation profiles confirm this decrease in WSS for ALE cases when compared to IB analogs (**Fig. 5**). Our single time-point comparison revealed comparable results, showing significantly lower WSS measures in ALE cases than in IB cases at both the inlet and bifurcation (**Table 3**). On average, IB WSS measures at the ICA inlet and bifurcation were 1.25 Pa (*p* < 0.0001) and 4.38 Pa (*p* < 0.0001) higher, respectively, than ALE values (**Table 4**). As was the case in stress and strain analyses, we continue to report the impact of vessel geometry on solver choice —finding that differences between IB and ALE WSS were more pronounced at the bifurcation than under inlet condition (*p* < 0.0001, **Table 4**).

**Fig. 7:**
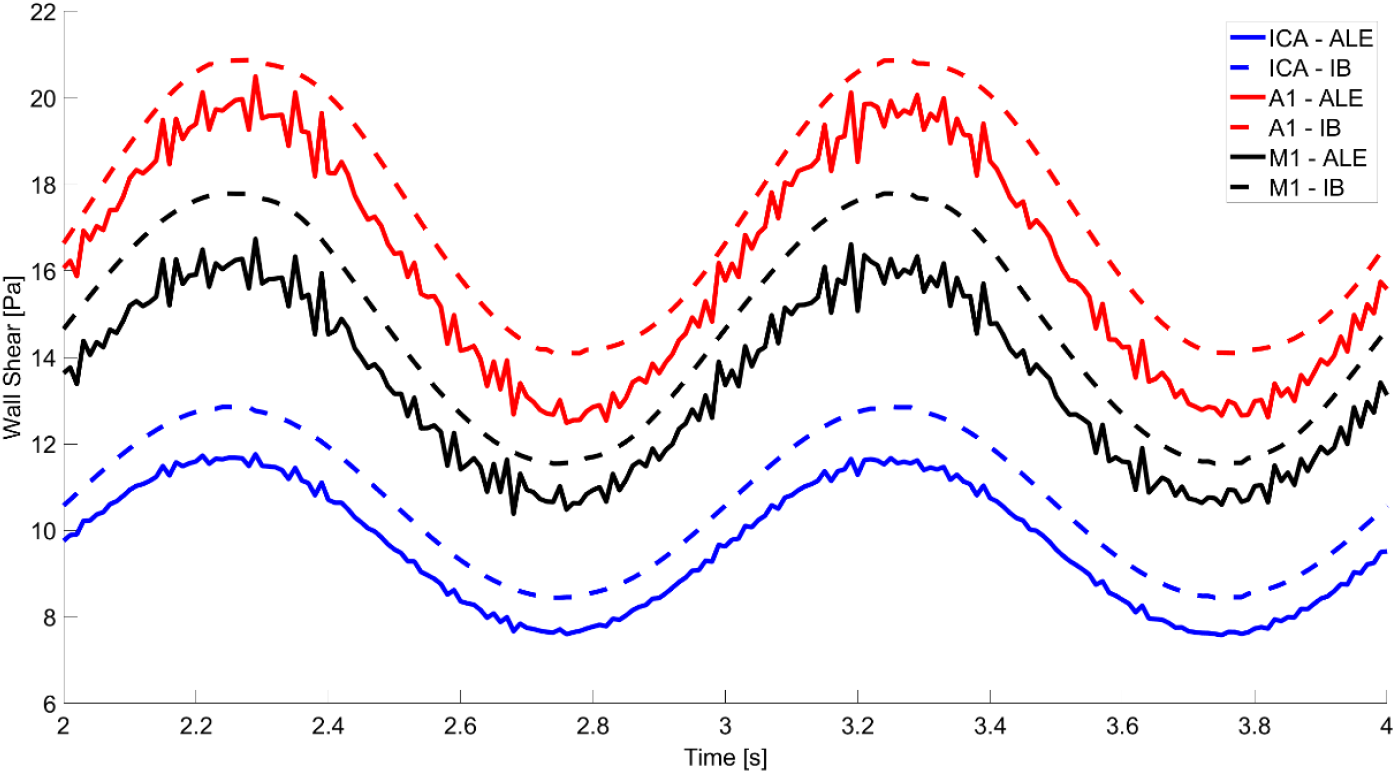
WSS comparison between IB and ALE methods at ICA bifurcation inlet and outlet geometries (A1 and M1).

### IV. Computation Time

As predicted by the inherent increase in simulation complexity with iterative solvers, the ALE-case computation required substantially more time to complete. While a 4 second, 400 step IB simulation converged to a solution in only 190 hours, an ALE simulation of the same task reached completion in 791 hours. In the monolithic case, Fluent simulations ran for 4 of the 190 hours, while the remaining computational time was spent solving viscoelastic responses in Mechanical.

## DISCUSSION

Our study confirms the feasibility of FSI modeling for the hemodynamic simulation of human cerebrovasculature, as proxied by our comparison simulation-derived flow velocity recordings with *in vivo* sonography data. Though simulation values appear elevated, this nominal variance is due to heightened flow velocities developed within the artificial entrance length preceding the ICA terminus. Future simulations may account for geometric differences between the entrance length and vessel conduit by adjusting inlet flow parameters.

In pursuit of a simulation standard, we provide a comparison between IB and ALE solvers for the FSI study of cerebrovascular tissue. We found that, despite increased computational demand, differences in simulation results warrant the use of ALE iterative formulation over the IB method for the modeling of cerebral vessels. Specifically, we report differences in flow velocity via qualitative analysis of streamline profiles and a quantitative description of velocity measurements. Velocities were notably lower when accounting for Mechanical influence on the Fluent domain. Velocity reductions are likely the result of an ALE iterative solver allowing the vessel wall motion to absorb a portion of fluid momentum. We also found deviations in strain, stress, and WSS between solving methods, suggesting that IB formulation fails to accurately capture a nuanced description of fluid motion and the behavior of the surrounding structure. One could, before, posit that IB simulation would be sufficient for the FSI modeling of cerebrovascular, as wall motion is relatively low compared to larger vessels of the periphery and previous modeling attempts of aortic valves [16]. However, our analysis refutes this notion and provides strong evidence for the use of ALE formulation.

Though much work remains before rapid life-course simulation of patient vessels reaches a point of relevant feasibility, FSI modeling has shown great promise in the field of vascular medicine by offering insights into the mechanical and hemodynamic characteristics of vessel tissue that may contribute to severe, acute conditions such as aneurysm rupture [2, 10, 17]. A number of *in silico* measures have already been identified to provide insight into the longitudinal integrity of vascular pathologies. WSS, in particular, was shown to predict rupture in an aneurysm model. In conjunction with our finding that monolithic formulation differs significantly from the ALE case in measures of WSS, this further strengthens the argument for partitioned formulation [10]. Interestingly, both circumstances of reduced and heightened WSS were identified as predictors of aneurysm rupture [18–20]; though a potentially controversial finding, this illustrates the necessity of accurate modeling. The ALE case allows for a higher degree of scrutiny over the behavior at each simulation node and is better suited to detect mechanical changes along a given vessel.

Regarding computation time, in this case, the ALE simulation took a little more than four times longer than the same simulation using IB. Running simulations on more cores with higher RAM, such as a cluster environment, would serve to decrease the computational time and reinforce our claim for ALE as the superior method, as ALE is always expected to be more intensive due to the additional data transfer and interaction between solvers. The evident computational cost with our current hardware precludes this current iteration of FSI from relevance as a diagnostic tool, which would require a significantly faster turnaround time; however, with a larger array of simulations of disparate patient geometries, a directory could be made to assist in guiding treatments.

### Future Directions

Cerebral blood vessels are known to possess non-linear viscoelastic character, but the degree to which this impacts the efficacy of an FSI model is unknown. It is entirely possible that these non-linear biomechanical properties are negligible for the purposes of a working small-vessel model and would only serve to add unnecessary computational load when a linear viscoelastic profile would suffice. Prior works have found that a non-linear, sigmoid model best predicted the viscoelastic response of large vessels at physiologic pressures, while a linear viscoelastic (Kelvin) model was sufficient for the consideration of relatively smaller vessels [21]. In smaller systems, the loading force elicits reduced non-linear stiffening at elevated pressures—despite being more rigid in a zero-stress state. It should also be noted that discontinuities in elastin and collagen content across vessel types contribute to this difference in elastic response, warranting further investigation of vascular histology for the improvement of future FSI models [21]. However, this prior comparison examined vessels from the aortic and carotid regions and warrants further investigation with the smaller-caliber vessels included in our study.

In addition, prior works have further postulated that physical restriction of small vessels by cerebrospinal fluid, surrounding astrocytes, or brain matter itself truncates vessel dilation amplitude [22]. Thus, restrictive interactions between the vessel and surrounding environment may impact FSIs within the vessel. The elastic modulus of brain tissue, glial cells, and CSF have already been characterized, but their potentially restrictive effects on vessel wall boundary conditions have yet to be elucidated [23–25]. As such, our future aims include an analysis of external boundary conditions by adding further constraints or pressures to the outer wall of the vessel.

### Limitations

This study demonstrates that quantifiable differences exist in the results of IB and ALE FSI simulation of flow through cerebrovascular tissue; nonetheless, there are a number of limitations to note. As discussed, this study did not impart boundary conditions on the vessel to proxy surrounding brain tissue and cerebrospinal fluid. It is possible that addition of an external pressure alters the simulation results by restricting vessel motion, however unlikely considering the limited wall displacement observed in this study. Second, we include no consideration for the different layers that comprise *in vivo* tissue. Instead of a 6 mm, uniform mesh layer, future studies could further delineate the Mechanical mesh with the addition of an internal elastic lamina, smooth muscle, and adventitial layers. Finally, the model parameters used in this study were not specifically derived from the patient whose vessel was modeled in angiography, but average values from the literature and the inflation testing of vessels from different donors (unpublished).

## REFERENCES

1. Hirschhorn M, Tchantchaleishvili V, Stevens R, Rossano J, Throckmorton A. Fluid-structure interaction modeling in cardiovascular medicine - A systematic review 2017-2019. Med Eng Phys. 2020, 78:1–13; doi:10.1016/j.medengphy.2020.01.008

2. Thiyagarajah N, Achey R, Rashidi M, Moore NZ. Computational fluid–structure interactions in the human cerebrovascular system: part 1—A review of the current understanding of cerebrovascular biomechanics. ASME J of Medical Diagnostics. 2022, 5; doi:10.1115/1.4053943

3. Shim JJ, Maas SA, Weiss JA, Ateshian GA. A Formulation for Fluid Structure-Interactions in FEBio Using Mixture Theory. J Biomech Eng. 2019; doi:10.1115/1.4043031

4. Moerman KM, Konduri P, Fereidoonnezhad B, Marquering H, van der Lugt A, Luraghi G, et al. Development of a patient-specific cerebral vasculature fluid-structure-interaction model. J Biomech. 2022, 133:110896; doi:10.1016/j.jbiomech.2021.110896

5. Byun HS, Rhee K. CFD modeling of blood flow following coil embolization of aneurysms. Med Eng Phys. 2004, 26:755–761; doi:10.1016/j.medengphy.2004.06.008

6. Cha KS, Balaras E, Lieber BB, Sadasivan C, Wakhloo AK. Modeling the interaction of coils with the local blood flow after coil embolization of intracranial aneurysms. J Biomech Eng. 2007, 129:873–879; doi:10.1115/1.2800773

7. Valencia A, Ledermann D, Rivera R, Bravo E, Galvez M. Blood flow dynamics and fluid–structure interaction in patient-specific bifurcating cerebral aneurysms. Int J Numer Methods Fluids. 2008, 58:1081–1100; doi:10.1002/fld.1786

8. Torii R, Oshima M, Kobayashi T, Takagi K, Tezduyar TE. Fluid–structure interaction modeling of a patient-specific cerebral aneurysm: influence of structural modeling. Comput Mech. 2008, 43:151–159; doi:10.1007/s00466-008-0325-8

9. Taylor CA, Hughes TJR, Zarins CK. Finite element modeling of blood flow in arteries. Comput Methods Appl Mech Eng. 1998, 158:155–196; doi:10.1016/S0045-7825(98)80008-X

10. Achey R, Thiyagarajah N, Rashidi K, Rashidi M, Moore NZ. Computational fluid– structure interactions in the human cerebrovascular system: part 2—A review of current applications of computational fluid dynamics and structural mechanics in cerebrovascular pathophysiology. ASME J of Medical Diagnostics. 2022, 5; doi:10.1115/1.4054124

11. Bazilevs Y, Hsu MC, Zhang Y, Wang W, Kvamsdal T, Hentschel S, et al. Computational vascular fluid-structure interaction: methodology and application to cerebral aneurysms. Biomech Model Mechanobiol. 2010, 9:481–498; doi:10.1007/s10237-010-0189-7

12. Souli M, Ouahsine A, Lewin L. ALE formulation for fluid–structure interaction problems. Comput Methods Appl Mech Eng. 2000, 190:659–675; doi:10.1016/S0045-7825(99)00432-6

13. Fukazawa K, Ishida F, Umeda Y, Miura Y, Shimosaka S, Matsushima S, et al. Using computational fluid dynamics analysis to characterize local hemodynamic features of middle cerebral artery aneurysm rupture points. World Neurosurg. 2015, 83:80–86; doi:10.1016/j.wneu.2013.02.012

14. Mount CA M, Das J. Cerebral Perfusion Pressure. StatPearls. Treasure Island (FL): StatPearls Publishing; 2018. https://www.ncbi.nlm.nih.gov/books/NBK537271/. Accessed 10 October 2024

15. Chen T. Determining a Prony Series for a Viscoelastic Material From Time Strain Data Varying. 2000. 20000052499.pdf (nasa.gov). Accessed 10 October 2024

16. Bavo AM, Rocatello G, Iannaccone F, Degroote J, Vierendeels J, Segers P. Fluid-Structure Interaction Simulation of Prosthetic Aortic Valves: Comparison between Immersed Boundary and Arbitrary Lagrangian-Eulerian Techniques for the Mesh Representation. PLoS ONE. 2016, 11:e0154517; doi:10.1371/journal.pone.0154517

17. Mut F, Löhner R, Chien A, Tateshima S, Viñuela F, Putman C, et al. Computational hemodynamics framework for the analysis of cerebral aneurysms. Int J Numer Method Biomed Eng. 2011, 27:822–839; doi:10.1002/cnm.1424

18. Chitanvis SM, Dewey M, Hademenos G, Powers WJ, Massoud TF. A nonlinear quasi-static model of intracranial aneurysms. Neurol Res. 1997, 19:489–496; doi:10.1080/01616412.1997.11740846

19. Xiang J, Natarajan SK, Tremmel M, Ma D, Mocco J, Hopkins LN, et al. Hemodynamic-morphologic discriminants for intracranial aneurysm rupture. Stroke. 2011, 42:144–152; doi:10.1161/STROKEAHA.110.592923

20. Cebral JR, Mut F, Weir J, Putman C. Quantitative characterization of the hemodynamic environment in ruptured and unruptured brain aneurysms. AJNR Am J Neuroradiol. 2011, 32:145–151; doi:10.3174/ajnr.A2419

21. Valdez-Jasso D, Bia D, Zócalo Y, Armentano RL, Haider MA, Olufsen MS. Linear and nonlinear viscoelastic modeling of aorta and carotid pressure-area dynamics under in vivo and ex vivo conditions. Ann Biomed Eng. 2011, 39:1438–1456; doi:10.1007/s10439-010-0236-7

22. Gao Y-R, Greene SE, Drew PJ. Mechanical restriction of intracortical vessel dilation by brain tissue sculpts the hemodynamic response. Neuroimage. 2015, 115:162–176; doi:10.1016/j.neuroimage.2015.04.054

23. Zhang C, Liu C, Zhao H. Mechanical properties of brain tissue based on microstructure. J Mech Behav Biomed Mater. 2022, 126:104924; doi:10.1016/j.jmbbm.2021.104924

24. Lu Y-B, Franze K, Seifert G, Steinhäuser C, Kirchhoff F, Wolburg H, et al. Viscoelastic properties of individual glial cells and neurons in the CNS. Proc Natl Acad Sci USA. 2006, 103:17759–17764; doi:10.1073/pnas.0606150103

25. Jurjević I, Rados M, Oresković J, Prijić R, Tvrdeić A, Klarica M. Physical characteristics in the new model of the cerebrospinal fluid system. Coll Antropol. 2011, 35 Suppl 1: 51–56; https://urn.nsk.hr/urn:nbn:hr:105:047138

26. Nagai Y, Kemper MK, Earley CJ, Metter EJ. Blood-flow velocities and their relationships in carotid and middle cerebral arteries. Ultrasound Med Biol. 1998, 24:1131–1136; doi:10.1016/s0301-5629(98)00092-1

27. Ringelstein EB, Kahlscheuer B, Niggemeyer E, Otis SM. Transcranial Doppler sonography: anatomical landmarks and normal velocity values. Ultrasound Med Biol. 1990, 16:745–761; doi:10.1016/0301-5629(90)90039-f

